# Primate ACC encodes natural vocal interactions in a ‘cocktail party’

**DOI:** 10.1101/2025.10.17.683014

**Authors:** Arthur Lefevre, Vikram Pal Singh, Timothy J. Tyree, Jingwen Li, Jean-René Duhamel, Cory T. Miller

## Abstract

The Cocktail Party Problem (CPP)—extracting meaningful signals amid competing voices—remains poorly understood at the neural level, particularly in real-world contexts where it emerges naturally. We investigated the role of the anterior cingulate cortex (ACC), a structure implicated in social monitoring but rarely examined in relation to audition, to resolve the CPP in freely-moving marmoset monkeys engaged in ecological vocal exchanges. Analyses revealed that ACC is seemingly integral to resolving the CPP. Not only did neurons encode the calls of either the conversational partner or background callers, but this selectivity persisted even with overlapping background sounds, a hallmark of the CPP. Moreover, ACC activity reflected the conversational dynamics by encoding turn-taking structure, highlighting the importance of CPP resolution for navigating social interactions in complex natural soundscapes.

## Main Text

The Cocktail Party Problem (CPP) — the perceptual phenomenon by which we extract meaningful sounds from noise - is arguably the most persistent real-world challenge for acoustic communication systems, such as speech^1^. Despite its fundamental importance and decades of research, the neural mechanisms underlying the CPP remain elusive, particularly in human and nonhuman primates^2^. This gap reflects at least the three limitations in prior research.

First, most neurobiological research related to the CPP has been performed in highly controlled laboratory settings that minimize uncertainty, competition, and social contingency^1–3^. Under these conditions, resolving the CPP is reduced to a sensory discrimination problem, placing limited demands on neural systems involved in adaptive control or interaction monitoring. Notably, this context contrasts considerably with the acoustically dynamic, naturalistic environments where the CPP was first described^3^ and naturally arises. Second, CPP research has focused primarily on neocortical regions associated with human speech processing^2,4–6^, reflecting a speech-centric and phylogenetically narrow view of the problem. This emphasis obscures the fact that the CPP predates language and the expansion of these cortical circuits in primate evolution, suggesting that more conserved neural systems likely contribute to its resolution.

Third, and perhaps most critically, beyond the choice of experimental setting, the CPP has been conceptualized primarily as an auditory computation. In real-world social interactions, however, the problem is not simply identifying a sound source, but determining which signal is relevant, when to respond, and how that response fits within an ongoing exchange—demands that require tracking social context and interaction history rather than acoustic features alone. Such functions are not central features of the canonical auditory pathways^7^, but are hallmarks of brain regions involved in cognitive control and social monitoring.

Together, these gaps motivate consideration of brain regions traditionally outside speech and auditory processing networks. One such candidate is the anterior cingulate cortex (ACC), which has been largely ignored in studies of CPP despite its established role in vocal production^8^, vocal perception^9^, and, critically, social monitoring during interactions with conspecifics^10^. Because resolving the CPP enables individuals to communicate with conspecifics in complex natural environments^11,12^, we posited that uncovering its neural basis likely requires its study under similarly dynamic conditions^13^.

To address this issue, we leveraged the rich vocal communication system of marmoset monkeys^14^ — highly vocal New World primates^15,16^. Marmosets engage in a range of context-dependent vocal exchanges, including turn taking with different call types^12^, and produce vocalizations that are modulated by social dynamics^17–19^. Moreover, these primates thrive in acoustically cluttered environments, making them an ideal species for studying the neurobiology of the CPP under real-world conditions.

We hypothesized that the ACC will exhibit at least the three following mechanisms integral to resolving the CPP in natural communication scenes: (i) selectivity for call type during both vocal production and perception as well as social context, (ii) respond invariantly to vocalizations regardless of background sounds and (iii) encode structure of active vocal turn taking exchanges. To test these hypotheses, we recorded the activity of individual ACC neurons in freely-moving, naturally behaving marmosets in their home cage within the acoustically rich colony room (Fig 1A), enabling detailed analysis of neural responses during natural conversations with their cage-mate partner or with members of the colony amidst ongoing background vocalizations from other conspecifics.

**Fig. 1.**
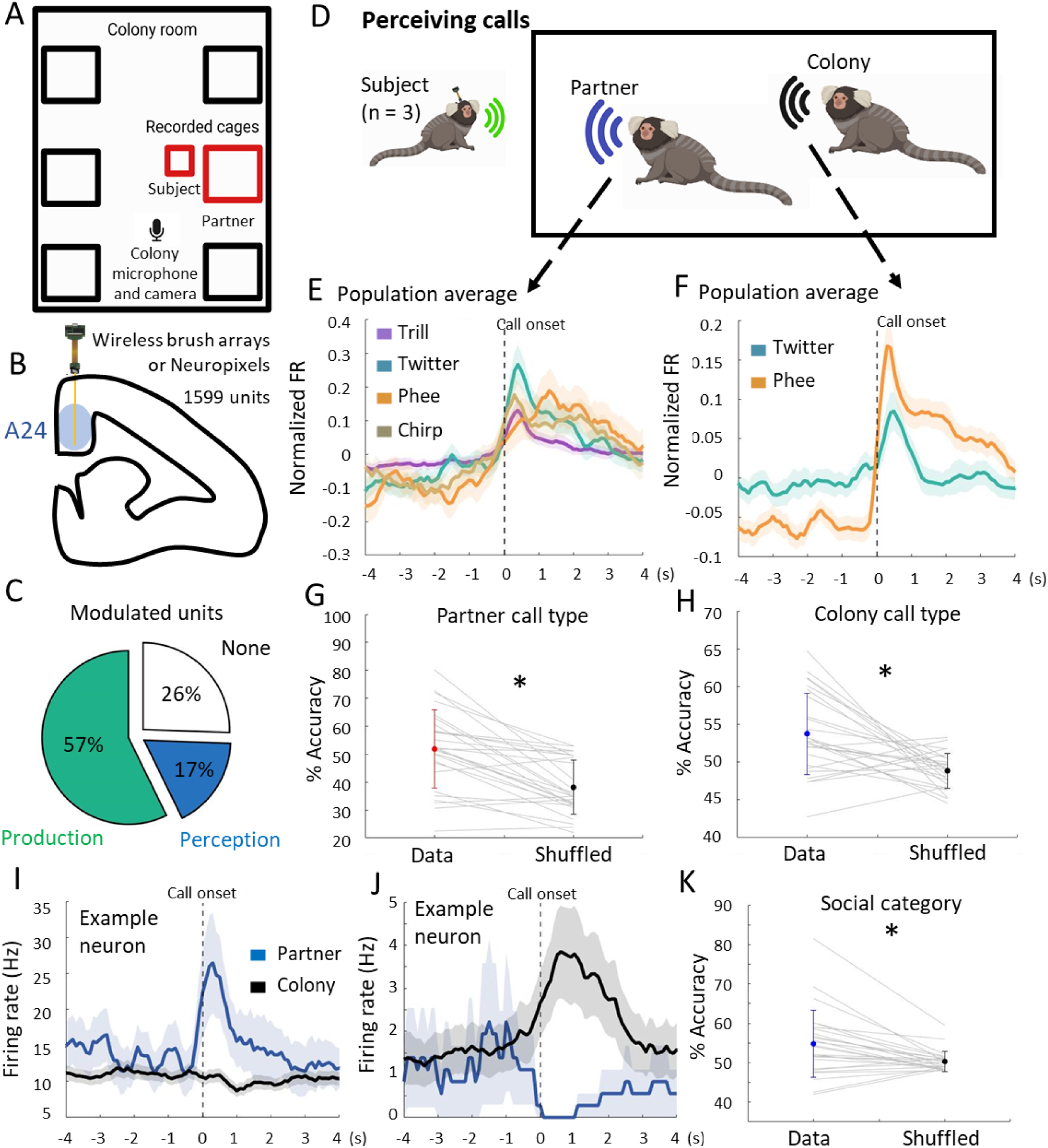
Segregated ACC neuronal populations encode vocalizations perceived from the partner and the colony. **A)** Schema of the experimental paradigm. *Top* – recordings took place directly in the home cage and transfer box in the colony room. *Center -* both subject and partner monkeys were equipped with a portable microphone, and the implanted subject was also equipped with a neurologger. *Bottom* – Neurons were recorded with brush arrays or Neuropixel probes implanted in ACC area 24. **B)** Coronal plane (AP = +12mm) at which probes were implanted. **C)** Percentage of neurons responding to vocalization production and perception. **D)** Vocalizations produced by the partner or other marmosets in the colony (vs produced ones) were analyzed separately. **R)** Normalized average increase from neurons with significantly upregulated activity during partner call perception. **F)** Normalized average increase from neurons with significantly upregulated activity during colony call perception. **G-H)** Percentage accuracy of the neuronal multiclass decoder for partner G) and colony (H) perceived call type for real data and data with shuffled labels. **I)** Example neuron responding specifically to twitter calls from the partner but not from the colony. **J)** Example neuron responding specifically to phee calls from the colony but not from the partner. **K)** Percentage accuracy of the neuronal SVM decoder for identity (partner vs colony) for real data and data with shuffled labels. *: p < 0.05, error bars indicate s.e.m, (standard error to the mean).

### Segregated ACC neuronal populations encode vocalizations perceived from the partner and the colony

To examine the neural basis of the CPP in marmosets, all experiments were performed in the noisy, colony room. The subject (2 females, 1 male) was placed in a small transparent box in front of their home cage that contained the partner (Fig. 1A). Using wearable microphones and an automated labeling algorithm^20^, analyses distinguished whether each call was produced by the subject, their partner, or the colony. We recorded ACC area 24 wirelessly using micro wire brush arrays or Neuropixels probes (Fig. 1B, Supplementary Fig. 1), yielding 1599 neurons and 38559 vocalizations across 46 sessions (Fig. 1B, Supplementary Fig. 2).

Similar to previous work^8,21–24^, we observed that ACC neurons were predominantly motor modulated, though a population were responsive to hearing the calls of conspecifics (i.e vocalization perception) (Fig. 1C). While ACC exhibits sensory responses to passive presentations of vocalizations^9,25^, whether this reflects a general response to conspecific calls or neurons in this substrate will display a selectivity to different call types necessary to resolve the CPP during natural vocal communication contexts was not clear. To test this, we first analyzed ACC neurons activity in response to hearing their partner’s vocalizations and those produced by other conspecifics in the colony (Fig. 1D). Analyses revealed that neurons in area 24 responded robustly to both partner calls (Wilcoxon signed rank tests, p < 0.05, n = 167, Fig. 1E, Supplementary Fig. 3C) and to colony calls (Wilcoxon signed rank tests, p < 0.05, n = 77, Fig. 1F). The majority of significantly modulated neurons responded to only a single call type (86.8%, Supplementary Fig. 3D). Given this degree of call type specificity in the activity of individual ACC neurons, we investigated whether call types were robustly encoded at the population level. A multiclass decoder, trained on neural activity of all neurons from 0 sec to +3 sec from call onset, revealed that individual call types from both the partner (Wilcoxon signed rank test, z = 4.16, p < 0.05) (Fig. 1G, Supplementary Fig. 4) and the colony (Wilcoxon signed rank test, z = 3.19, p < 0.05) (Fig. 1H, Supplementary Fig. 4) could be significantly decoded from ACC population activity.

We next examined whether ACC area 24 neurons separately encode different types of callers in the scene, namely partner calls and those produced by other marmosets in the colony. In other words, do these neurons simply encode call type or a combination of call type and social categories? We observed two distinct populations of neurons, one that responded only when hearing partner calls (Figure 1I) and the second that responded only to hearing other callers in the colony (Figure 1J). This suggests that ACC neurons parse conspecific calls into two categories - partner and others (i.e. colony) – and encode both. While these neural populations were almost entirely segregated; 2 neurons (out of 244) responded to both partner and colony calls, suggesting the existence of a minority of neurons with general auditory responses. Moreover, the selectivity was not biased to partner calls, with similar proportions of neurons encoding both categories of callers within the same ACC substrate (5.3% of neurons encoding partner phees or twitters, 4.8% encoding colony phees or twitters). At the very least, these neurons suggest that figure/ground separation of highly meaningful signals^1,3,26^ is encoded in ACC area 24, a foundational mechanism for resolving the CPP in natural acoustic scenes. To further validate this idea at the population level, we used neural population activity to decode whether a call had been produced by the partner or the colony using a support vector machine (SVM) classifier. Analyses revealed significant decoding of these social categories (Wilcoxon signed rank test, z = 2.91, p < 0.05) (Fig. 1K).

### ACC neurons respond invariantly to calls with and without overlapping vocalizations

To fully resolve the CPP, neural mechanisms that parse meaningful signals (such as partner’s voice) from acoustically overlapping background sounds are necessary^1,3^, particularly in the presence of similar vocalizations produced by conspecifics. To more closely examine how ACC responses were affected by background sounds, we investigated how neurons responded to vocalizations in the two following contexts^26^: ‘Simultaneous calls’ (i.e. calls that occurred concurrently with a colony call (Fig. 2A)) and ‘“Sequential calls” (i.e. calls that had no acoustic overlap with other vocalizations (Fig. 2B)^26^). Analyses focused on twitter calls because it had the largest population of perceptual neurons selective to partner calls (n = 54 neurons), thereby allowing comparison in both contexts.

**Fig. 2.**
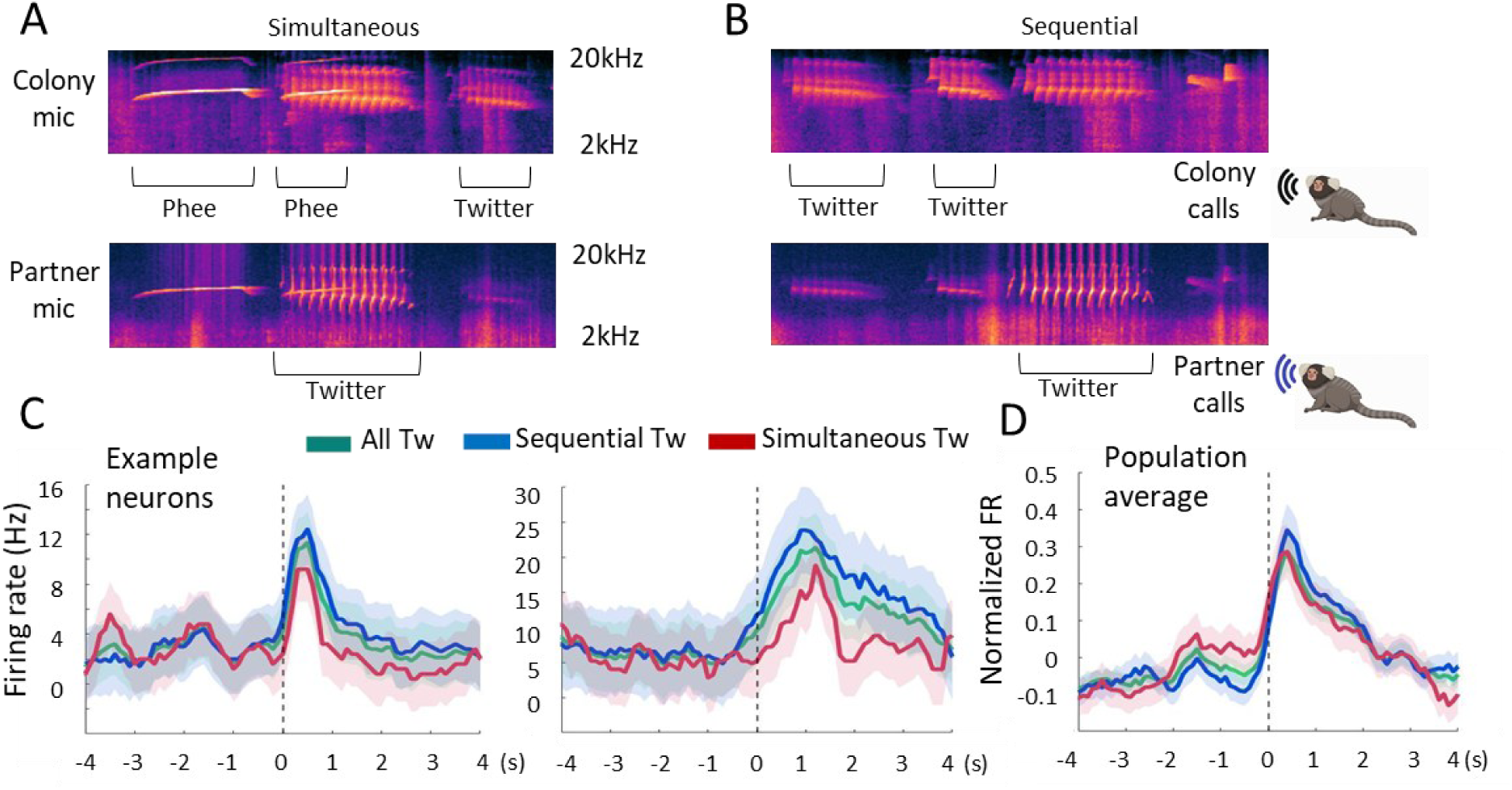
ACC neurons respond invariantly to calls with and without overlapping vocalizations. **A)** Example spectrograms from the colony and partner microphones showing the occurrence of a call from the partner (twitter) that is perceived simultaneously with a call from the colony (phee). **B)** Example spectrograms from the colony and partner microphones showing the occurrence of a call from the partner (twitter) that is perceived sequentially to calls from the colony (twitters). **C)** Example neurons responding to twitter calls from the partner that are unaffected by the presence (or absence) of simultaneous calls from the colony. **D)** Normalized average increase from neurons with significantly upregulated activity during partner twitter call perception is unaffected by the presence (or absence) of simultaneous calls from the colony. *: p < 0.05, error bars indicate s.e.m.

Analyses revealed that although neurons were strongly driven to partner twitter calls (Wilcoxon signed rank tests, p < 0.05), there was no statistical difference in responses between the sequential and simultaneous contexts (p > 0.05) (Fig. 2C, D), indicating that background sounds do not affect the responses of these neurons. In other words, single neurons in marmoset ACC area 24 selectively parse the meaningful signal – vocalizations produced by the partner - from overlapping background sounds (e.g., all other conspecific vocalizations), thereby exhibiting a key mechanism necessary to resolve the CPP. Following this logic, populations of neurons encoding partner calls and colony calls were almost entirely distinct (Fig. 1J, K). This suggests that attentional mechanisms in ACC^27–29^ enable neurons in this substrate to encode distinct calls produced by conspecifics that are temporally simultaneous.

### Marmosets Monitor Social Scenes to Optimize Vocal Turn Taking in a Cocktail Party

Resolving the CPP is essential for vocal communication in both human and nonhuman animals, where the ability to parse, produce, and respond to vocal signals amidst background noise enables meaningful interactions. To better understand how marmosets communicate in such environments, we examined vocal turn taking with different vocalizations in the noisy colony environment, a natural behavior observed in many animals including humans and marmosets^12^.

To quantify marmoset turn-taking in this contexts, we focused analyses on two distinct vocalization types — trills^14,20^ and twitters^14^ (Fig. 3A, B) — by aligning the partner and colony call rates on the subject’s vocalizations (Fig. 3C, D). Trills are low-amplitude calls exchanged between affiliated conspecifics in close proximity^14,20^, whereas twitters are composed of multiple sharply rising notes, mid-distance contact calls, with less affiliative component^14,30^. This natural ethological difference between these call types during turn-taking provided an opportunity to investigate the behavioral strategies marmosets use to communicate in noisy acoustic scenes. We found that partners produced significantly more trills in close temporal proximity to the subject’s trill onset (sliding Wilcoxon’s signed rank tests, 0.1 to 1.4s, p < 0.01, Fig. 3C) but no such relation was found for twitters between partners (Fig. 3C). Moreover, subjects produced the low amplitude trill calls when the colony twitter call rate was significantly lower than average (−1.5 to 0.6s, p < 0.01, Fig. 3D), effectively avoiding noisy periods in the colony. Finally, we found increase in colony twitter rate before and after subjects’ twitter meaning subjects were twittering in synchrony with the colony (1.5 to 2.5s), while avoiding interrupting each other, as represented by lower colony twitter rate at subjects’ twitter call onset (0.5 to 1.6s, p < 0.01, Fig. 3D). This pattern suggests that marmosets monitor the colony’s sound level and adopt context-dependent strategies for turn-taking with each call type to maximize respective communication signaling efficacy.

**Fig. 3.**
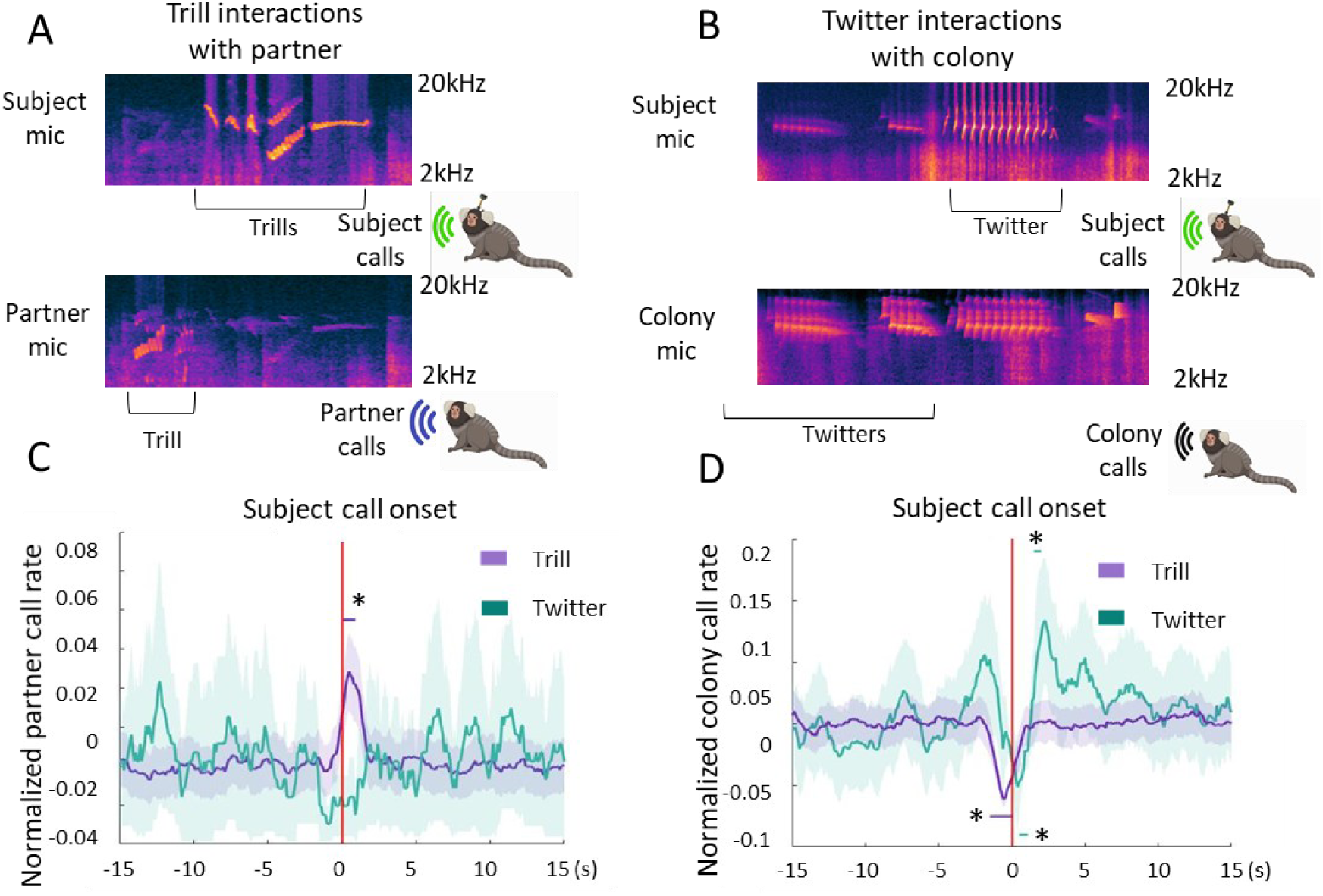
Marmosets Monitor Social Scenes to Optimize Vocal Turn Taking in a Cocktail Party. **A)** Example spectrograms showing trill to trill conversations between the subject and its partner. **B)** Example spectrograms showing twitter to twitter conversations between the subject and other monkeys in the colony. **C)** Normalized partner call rate for trills (purple) and twitters (orange) aligned on subject’s trills and twitters onsets. **B)** Colony call rate for twitters aligned on subject’s trills (purple) and twitters (orange).

### ACC neurons encode active turn taking with the partner and the colony

Although the CPP has been considered mainly as a perceptual problem, it is integral to communication in noisy socio-acoustic environments. Thus, in the present paradigm, our subjects were not only perceiving calls but also producing them, reflecting active communication. We analyzed neural activity during vocal production. Results showed that neurons exhibited a significant increased (n = 311, 19.4%) or decreased (n = 652, 40.8%) activity when the subject was producing a call (Wilcoxon signed rank tests, p < 0.05) (Fig. 4A-C, Supplementary Fig. 3A, B, D). Like call perception (Fig. 1F, G), the majority of neurons were highly selective to only one call type (83.3%, Supplementary Fig. 3D). However, a subset of neurons exhibited responses to multiple types of vocalizations (Supplementary Fig. 3D).

**Fig. 4.**
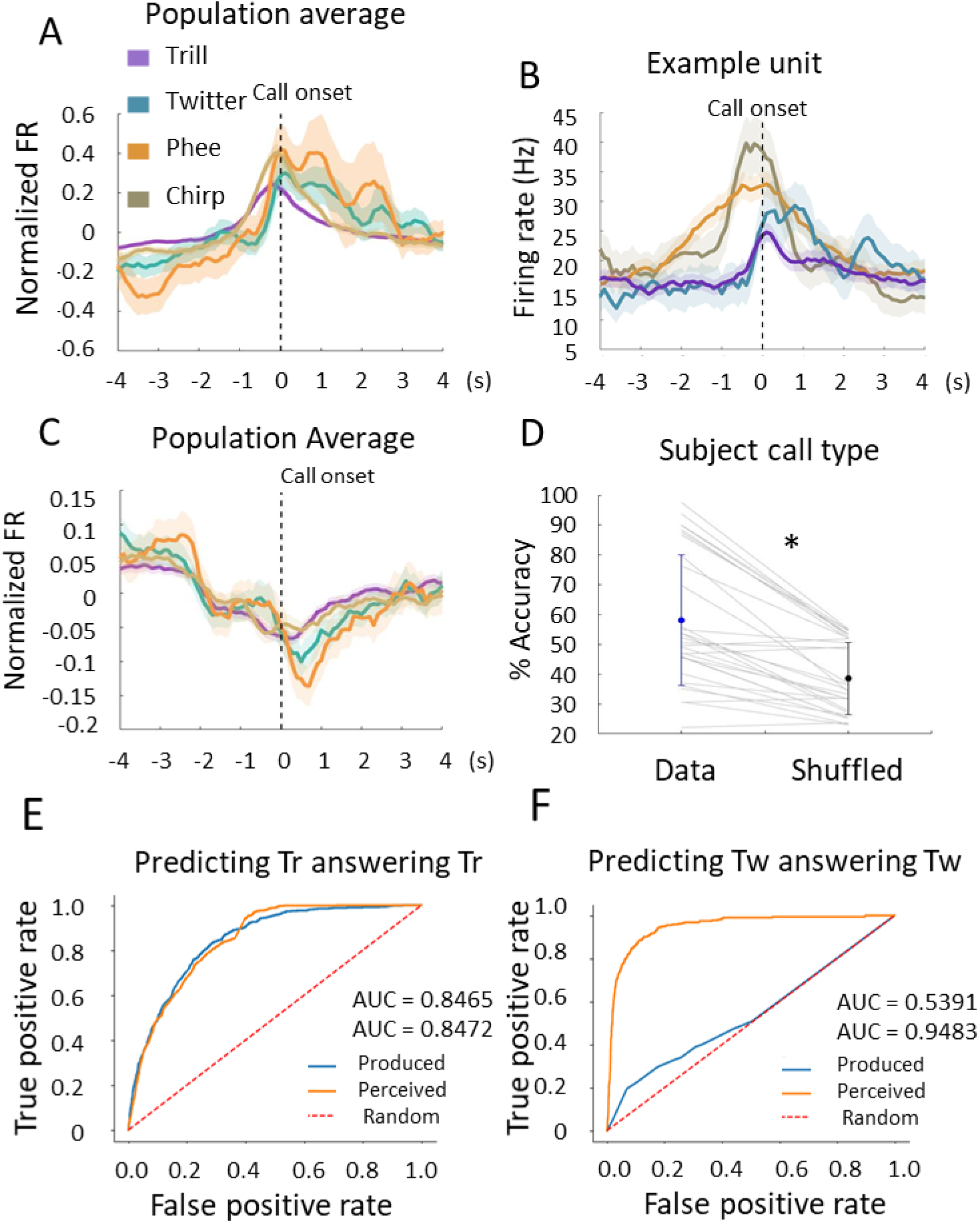
ACC neurons encode active turn taking with the partner and the colony. **A)** Normalized average increase from neurons with significantly upregulated activity during call production. **B)** Example neuron responding to the production of several call types. **C)** Normalized average decrease from neurons with significantly downregulated activity during call production. **D)** Percentage accuracy of the neuronal multiclass decoder for produced call type for real data and data with shuffled labels. **E-F)** Receiver Operator Characteristic (ROC) traces for our neural decoder trained on predictive time bins representing whether trills answered trills with the subject’s partner in panel E, the answering of twitter calls with the colony in panel F. Blue solid lines represent calls initially produced by the subject while orange solid lines represent calls initially perceived by the subject. Random chance is represented by the red dashed line as a guide. The area under the curve (AUC) quantifies the success of the neural decoder as our outcome measure. *: p < 0.05, error bars indicate s.e.m.

We quantified whether produced call type could be robustly decoded from neural population activity using a multiclass decoder trained on calls produced by subjects. Results showed that call types could be accurately classified based on ACC activity (Wilcoxon signed rank test, z = 4.54, p < 0.05) (Fig. 4D, Supplementary Fig. 5). This decoding was dependent on the number of neurons rather than on the ones significantly modulated by call production (Supplementary Fig. 5). Together, these data show that ACC area 24 neurons encode call production before, during and after call onset with a high degree of selectivity, consistent with results in other motor systems^31^.

To investigate whether ACC neurons are sensitive to the social context in which calls are produced, we employed a time sensitive decoding analysis designed to identify periods with maximal differences in neural activity between different contexts (see Methods and ^32^ for details). Specifically, we applied this approach to test whether population activity could reliably classify whether a given call occurred as part of a structured turn-taking exchange or independent of these social interactions. Turn-taking was defined as an event in which subjects produced a call within 5 seconds (for partner trill calls) or 10 seconds (for colony twitter calls) of a conspecific producing the same call type (Fig. 3A, B).

Analyses revealed that the behavioral context – whether the vocalization occurred in a turn-taking exchange or not – could be accurately classified from ACC area 24 population activity. Trill calls produced (accuracy: 86.9%) or perceived (accuracy: 82.7%) (Fig. 4E, Supplementary Fig. 6C) were accurately classified as isolated or part of a turn-taking bout with the partner, according to one-sided Mann-Whitney’s U-tests (z = 2.48×10^5^, p < 0.0001, N_calls_ = 596 produced, N_calls_ = 776 perceived). Likewise, twitter calls produced (accuracy: 68.3%) or perceived (accuracy: 96.0%) (Fig. 4F, Supplementary Fig. 6D) were classified with significant statistical accuracy according to a one-sided Mann-Whitney’s U-test (produced: z = 3.1x10^4^, p = 0.0105, N_calls_ = 236; perceived: z = 8.5x10^4^, p < 0.0001, N_calls_ = 314).

These results indicate that ACC area 24 encodes not only the calls themselves, but information about social context. Because many of these turn-taking exchanges occurred during noisy acoustic periods, particularly for twitter calls, this also suggests that levels of background noise do not affect ACC activity during a coordinated social interaction, further emphasizing the role of this neural substrate to resolve the CPP.

## Discussion

Here, we investigated the role of ACC area 24 in resolving the CPP as freely-moving marmosets engaged in natural vocal communication in a noisy environment. We found that single neurons selectively responded to partner vocalizations or colony calls, effectively parsing different categories of conspecific speakers in the acoustic scene (Fig. 1). Critically, neurons selective for partner calls responded invariantly when calls overlapped acoustically with other sounds, showing that neural mechanisms to distinguish meaningful voices from other interfering voices, are present in marmoset ACC area 24 (Fig. 2). These findings suggests that ACC may be a keystone neural substrate for resolving the CPP, highlighting the significance of this process for mediating social interactions in natural scenes, rather than solely a challenge for audition.

As a midline forebrain area conserved across all mammals^33^ that is both anatomically connected to fronto-lateral^34^ and auditory cortex^35^, and modulated by attention^27–29^, it may perhaps not be surprising that the ACC supports resolution of the CPP. Although not classically considered part of the language network, results here show that ACC neurons are involved in a range of communicative functions, including representing each of the call types produced or perceived in vocal exchanges (Fig. 1, 3). Recent marmoset fMRI studies have indeed implicated ACC in vocalization processing^9,25^, suggesting strong interactions with the broader communication network. Moreover, if the CPP is viewed not as a problem of literal language comprehension but as one of identifying and tracking communicative partners in noisy social environments, resolving it requires listeners to recognize each speaker’s voice. Afferent connections from hippocampus, which robustly encodes identity in social signals^32^, to area 24^35^, could provide this necessary input. Taken together, these findings and the underlying anatomical connectivity suggest that ACC area 24 plays a more central role in resolving the CPP than previously thought, potentially emphasizing that the CPP is not solely a challenge for audition but one of sociality.

Marmosets engage in vocal turn-taking between two or more individuals ^12,14,36,37^. In the noisy colony setting, we observed differences in the relative timing of the call types produced in these social exchanges in the presence of background noise. Whereas mid-distance twitter calls were produced in turn-taking bouts with conspecifics in the colony, trill calls – a low amplitude call used when in close visual contact - were mostly produced during periods of silence in the colony. This shows that marmosets monitor the acoustic landscape to optimize signaling efficacy during communicative exchanges^38–41^. Moreover, ACC appears to play a role in this process, as the social context in which these calls were produced and perceived could be reliably decoded from neural activity alone (Fig. 4). In parallel, fMRI findings in humans point out that ACC is activated by listening to conversations rather than monologues^42^, and more generally that ACC key role in theory of mind is integral to natural language processing^43–46^, and language production, as ACC lesions in humans produce akinetic mutism^47^. Finally, ACC is involved in social monitoring^48^, making it a logical node to be recruited for processing the CPP. Because most studies in this line of research were performed in the visual modality^10^, it suggests that ACC integrates multisensory information to guide social behavior.

Growing evidence suggests broader roles for ACC in social beahviors^10,45,46^, functions that may be particularly pertinent for resolving the CPP during natural communication. One influential framework, the Foraging Value Theory (FVT)^49^, states that in a foraging for food context, the dACC is monitoring the value of the exploited option versus moving to explore other potential options. While originally developed in the context of food foraging, this framework may extend naturally to social domains. From this perspective, the marmosets in the present study can be viewed as foraging within a social environment, dynamically adjusting their vocal interactions to maximize social reward. In humans, answering and being answered can be intrinsically rewarding^50^. If similar reward contingencies apply to marmosets, area 24 may be optimizing the behavior to maximize social reward (and even perhaps in competition with other rewards/motivational factors). Consistent with this view, the tendency of marmosets to preferentially engage their partner when the colony is relatively quiet may reflect a reward-seeking strategy that complements, rather than replaces, purely acoustic optimization mechanisms. Conversely, periods of heightened colony activity may increase the relative value of interacting with other individuals, favoring broader social engagement. Because our neuronal recordings show an encoding of the social context (answers *versus* non-answers), area 24 function in this ecological context could be consistent with a social version of the FVT, although it should be noted that other ecological contexts might reveal additional functions of the area 24. More broadly, these findings illustrate how recording neural activity during spontaneous, naturalistic behavior can reveal functions of brain areas that remain obscured under constrained laboratory conditions.

By recording neural activity during natural vocal exchanges of freely-moving primates in rich acoustic environments, we discovered that the anterior cingulate cortex area 24—a region long overlooked in this domain—implements core computations necessary for resolving the CPP. Certainly, ACC is not solely responsible for resolving the CPP, our results here identify it as a central, integrative hub that coordinates the perceptual, contextual, and action-related processes necessary for effective communication in noisy real-world environments. Moreover, these results demonstrate that CPP resolution extends beyond sound parsing to organizing socially meaningful interactions, requiring the integration of perception, behavioral response, and social context. This work showcases the transformative potential of naturalistic neuroscience and points to a broader reevaluation of how and where the brain solves real-world cognitive challenges^13^.

## Acknowledgments

Funding

This work is supported by grants to AL (Marie Sklodowska-Curie fellowship 101018877 and European Research Council Starting grant 101116110) and to CTM (National Institute of Health R01 DC 012087).

## Author Contribution

Conceptualization: AL, CTM Formal Analysis: AL, TJT, JL

Methodology: AL, VPS, CTM Investigation: AL, VPS

Funding acquisition: AL, JRD, CTM

Project administration: AL, CTM

Supervision: JRD, CTM

Writing – original draft: AL, CTM

Writing – review & editing: AL, VPS, TJT, JL, JRD, CTM

## Data and material availability

All data and code are available on request to the corresponding author.

## Supplementary material

### Material and methods

#### Animals

Four adult marmosets (*Callithrix jacchus*) living in bonded pairs (1 female and 1 male per cage) for at least 3 months were used for this experiment. All pairs were housed in a room with other marmoset cages including two adjacent cages with visual separators (between 41 and 47 marmosets in total). Three of the marmosets, monkey S (female), G (female) and L (male) were implanted with electrodes while monkey K (male) was only used for behavior. Animals had unrestricted access to food and water. All experiments were approved by the UCSD Institutional Animal Care and Use Committee and were performed in the Cortical Systems and Behavior Laboratory at University of California San Diego (UCSD).

#### Behavioral paradigm

Marmosets were first habituated to be handled and taken out in transport boxes by the experimenters for a month. Once habituated to handling (measured by assessing the comfort of animals while being handled and taking treats from the experimenters’ hands), we gradually habituated them over one or two months to wear the leather harness containing the microphone until they spent less than 20% of the time chewing on it. Experimental sessions lasted between 1 hour to 1.5 hours. We recorded only one pair of monkeys per session. During a session, the pair was taken out of their home cage in transport boxes and brought to an adjacent experimental room for preparation. We then equipped both the subject and the partner monkey with the custom-made leather harness containing a microphone (micro voice recorder, Spycentre Security®). Additionally, we connected a neurologger to the subject’s head-cap. Both monkeys were brought back to the colony room. The partner was placed back in its home-cage and the subject was kept in a transparent transport box placed at the entrance of the home-cage (Fig 1A, Fig S2A) giving the pair visual and auditory access to each other. To record colony calls, we used an omnidirectional microphone (H3-VR, Zoom®) placed in the middle of the room. This paradigm allowed us to record acoustic interactions between the subject and its partner, and between the subject and the colony. In marmosets, which are capable of turn-taking, these interactions involve the coordinated exchange of vocal signals between individuals, characterized by social coordination and minimal acoustic overlap, allowing speakers to alternate the timing of their calls within a shared acoustic space.

#### Neurophysiological procedures

To target the desired brain regions for electrophysiological recording using external probes, the brain anatomy of the subjects was imaged using a 3T MRI (Siemens MAGNETOM Prisma) scanner and a human knee coil (Siemens/QED Transmit/Receive 15 channels). T2 and T1 weighted images of the whole marmoset brain were obtained in an anaesthetized animal placed in a custom non-magnetic 3D printed stereotax with earbars and bite plate. We adapted the coordinates from the Paxinos atlas^51^ of the marmoset brain based on individual brain size. We targeted the left hemisphere area 24 (Site 1: AP +12mm, ML +1mm, Site 2: AP +13.5mm, ML +1mm). On the day of surgery, monkeys were intubated and maintained under isoflurane anesthesia, then placed in a surgical stereotaxic frame (Kopf). We performed a skin incision and a craniotomy (1.5mm diameter), then inserted a ground electrode between the dura and the skull before covering the skull with metabond dental cement (C&B Metabond®). We employed 2 different strategies. For monkey S and G, we used micro wire brush arrays (Microprobe Maryland, USA, Monkey S:1x64 channels, Monkey G: 2x32 channels with 2mm space in between) with microdrives (custom made from^52^, for Monkey G, an additional hole was pierced and the drive adapted to hold 2 micro brush arrays). The electrode guide tube(s) (Microlumen, High performance medical tubing, 310-I.5 PTFE), containing a 24G needle, attached to the microdrive was/were inserted straight into the brain at a depth of 1.5mm. The needle(s) was/were retracted and the electrode(s) array(s) were lowered into the guide tube(s) until they protruded out of the guide tube by 1mm. Kwik Sil (World Precision Instrument) was used to fill the craniotomy and seal the gap at the top of the guide tube(s). Then the Microdrive and the omnetics connector were cemented to the metabond using dental acrylic, after soldering the ground electrode to the array ground wire. Finally, the skin was glued to the metabond using Vetbond (3M). For monkey L, we employed Spikegadget wireless neuropixel system. The procedure was similar, except that a durotomy was performed and the 2 probes (Npx 1.0 and Npx NHP 1.0 10mm) inserted 5mm deep from the cortical surface before we covered them with kiwk-Sil and fixed them directly to the skull with metabond. Probes were then connected to the Spikegadget head cap that was built around the electrodes and cemented to the metabond. After the surgery, monkeys received analgesic treatment for 3 days. Recording sessions started 2 weeks after the surgery for monkey S and G and were performed twice per week, and started 4 days after surgery and happened every day for monkey L (monkey S: 21 sessions, monkey G: 19 sessions, Monkey L: 6 sessions). Implantation sites were verified by structural MRI for monkey G and S (Fig S1B), but this could not be done for monkey L.

To wirelessly record neurons, we used a 64 channels Neurologger (Deuteron Technologies) for monkey S and G with a sample rate of 32 KHz, and Neuropixels Datalogger Headstage (SpikeGadgets) with a sample rate of 30 KHz for monkey L.

### Data preprocessing

#### Vocalizations

To automatically label subjects’ and partners’ vocalizations, we retrained the algorithm described in^20^ on the data recorded from our 4 monkeys with the wearable microphones. This artificial neural network^53^ identifies the call type and determine the source (subject or partner) by comparing the amplitude. If the amplitude is equal in both microphones, the call is considered to be coming from the colony. Following this step, all audio files were manually corrected to remove eventual mislabeling by the algorithm and determine precisely (<20ms) each call onset and offset. We considered only 4 call types: trill, twitter, phee and chirp. Other calls (tsik, ek, chatter, trillPhee, whistle) had too few occurrences to be analyzed. Additionally, to investigate the cocktail party effect, Twitter calls coming from the partner were labeled as “simultaneous” or “sequential” depending on if a call from the colony was overlapping or not with it.

For calls coming from the colony (recorded with the omnidirectional microphone), a first pass was done with ACDC neural network (https://github.com/mineraldragon/ACDC_2022) and then corrected manually. We only labeled Twitter and Phee calls from the colony, as other call types such as trill and chirp were not loud enough to be recorded consistently, or had too few occurrences of them (eg tsik, chatter, etc.). Note that for technical reasons, colony calls were not always recorded (34 out of 46 sessions with colony recordings).

We removed all calls that were happening less than 2 seconds after another call from the same source, to ensure that neuronal responses analyzed were not influenced by a recent prior call.

This was the case for all analyses except for PSTH of phee calls perceived from the partner (Fig 1F), due to low number of calls. Thus, Fig 1F phee call data may contain residual activity from successive calls.

Finally, we removed calls that occurred when the neural signal was poor due to artifacts, defined as calls for which 20% of the neural signal (from 1sec before to 1 sec after the call onset) was labeled as artifact (see neural data).

#### Neural data

All neural data were band pass filtered (300-7000Hz). Signal from brush arrays (but not Neuropixel) contained artifacts, that were removed by setting signal value to 0 if the standard deviation of one channel was above a threshold (arbitrarily set to 100 after visual inspection of signal). If more than 20% of a session was removed, the session was not included in our analyses. We then used Kilosort 2^54^ to performed automatically the spike sorting for both micro brush arrays and Neuropixels data. Each identified unit was manually checked and validated and labeled as single or multi-unit or rejected based on the waveform shape and auto-correlogram using Phy (open-source Python library for spike visualization and curation). Only units with a minimum firing rate of 0.5Hz were considered. Across the 3 monkeys (Monkey S: units, Monkey G: 574 units, Monkey L: 794 units) a total of 1599 units were identified, of which 932 were well isolated single units (using the PCA). However, because we did not see differences between single and multi-units in later analyses, we decided to pull them together and to refer to them (single and multi-neurons) as neurons throughout this article. All PSTH in this article are showing results from well isolated single units.

#### Synchronization

All data streams (3 microphones and neurologger) were synchronized using 250ms audio tone pulses played via a buzzer generated by an ESP32 microcontroller programmed using Arduino IDE.

All .wave files were manually aligned with Audacity® (v 3.4). Two tones (one at the start of the session and another at the end) were used to bookend the whole session which allowed us to identify and correct the temporal drift of each microphone in Audacity using the “change speed” functionality. Audacity was able to correct the temporal drift in each microphone to less than 20ms. Following this, a MATLAB script was used to randomly remove data points until the temporal time drift was reduced to <1ms. This did not influence the labeling of calls by the artificial neural network.

A 100 mv TTL pulse (100 ms) was sent to the Neurologger using the same ESP32 microcontroller neuron at the beginning of the session. The time difference between the Neurologger TTL and first acoustic pulse was used to align the timing of vocalizations onset and offset to the neurologger time. Using this approach, we were able to synchronize different systems with varying sampling frequencies to a common master clock.

### Statistical analyses

<u>PSTH responses:</u> Firing rate was computed over 100ms bins. To test neurons’ response to produced or perceived calls we first employed a peak detection approach. For each call type, if there were at least 5 calls, we calculated the mean firing rate between -5s to -2s before call onset for produced calls and between -5sec to 0sec before call onset for perceived calls as a baseline. We then searched for the peak response between -2s and +1s around call onset for produced calls and between 0sec and 3sec after call onset for calls perceived. To ensure a minimum peak width, we then tested if the bins around the peak bin reached at least 50% of the peak value (threshold = baseline +- 0.5*(peak-baseline)) to have at least 3 consecutive bins (300ms) beyond this threshold. Finally, we tested if the peak values were significantly higher than the baseline values using the non-parametric paired Wilcoxon signed rank test. We used this approach to allow for time flexibility because we observed that neurons responded at various time windows (Fig. 4, Supplementary Fig. 6). Note that both maximum and minimum peak responses were computed to look for excited and inhibited neurons. Each neuron was tested for each produced and perceived call types.

Finally, the PSTH traces reported in Fig. S6 were first computed at a bin width of 0.3 seconds, then up-sampled to a bin width of 0.1 seconds using linear interpolation that assigned knots to the mean time of any given bin, and finally smoothed with a sliding window with a duration of 2 seconds using a Savitzky-Golay filtration. Uncertainty of these demonstrational PSTH traces indicates 95% confidence estimated via bootstrap and is shown by the relatively horizontal, yet curved shaded regions in Supplementary Fig. 6.

<u>Decoding:</u> To decode call type or caller category, we employed a multiclass decoder (error-correcting output codes model, MATLAB function *fitcecoc*) with a one-versus-all coding design. For each session, the input was the average firing rate of each neuron between -2s and +1s around call onset for produced calls and between 0sec and 3sec after call onset for calls perceived. The number of calls of each type (trill, twitter, phee or chirp) or caller (partner or colony) was equalized by subsampling categories with more calls down to the category that had the least. Only calls that had at least 10 occurrences were considered, leading to different number of categories and therefore chance levels for each session (25%, 33% or 50%). The decoder was trained on 80% of the data and tested on the remaining 20%. To control for random effects, we also decoded on neural data with shuffled labels. This procedure was repeated 100 times to obtain a representative average decoding value for the data and the shuffle. Finally, we used Wilcoxon sign rank test to compare the average decoding accuracies obtained on the real and the shuffled data.

*<u>Time bins:</u>* To quantify the neural representations that identified whether a trill with the subject’s partner was answered or whether a twitter with the subject’s colony was answered, we used a predictive time bin analysis similar to previous work in the lab^32,55^. In this study, we applied the same four-fold stratified cross-validation both to detect predictive time bins and to decode their apparent firing rates using an ensemble of gradient-boosted decision trees^56^. Recording sessions were selected for these analyses if they had at least ten repeatedly spiking neurons recorded, at least four recorded calls that were answered, and at least four recorded calls that were not answered. A trill call was considered an answer if it happened within 5 seconds of a trill call from the other monkey, and a twitter call was considered an answer if it happened within 5 seconds of a twitter call from other colony monkeys. These timings were based on previous literature^20,57^. As a control, the same analysis was repeated with pseudo-randomly shuffled labels, and decoding results are reported in the main text and in Fig. 4. All time bins considered only the times from five seconds before to five seconds after call onset and were constrained to be no briefer than four hundred milliseconds in duration, resulting in a median activation time of t=-0.17 (IQR: -2.34 through +2.14) seconds (N_bins_ = 4326 predictive time bins) relative to call onset at t=0. Predictive time bins were also constrained to be non-overlapping for any given neuron such that no action potential was considered twice for a given type of neural representation. Reported tree-based decoder accuracies were computed using a threshold parameter value that maximized apparent accuracy. Area under the ROC curves provide an alternative outcome measure that is parameterless and is reported in the Fig. 4E, F and Supplementary Fig 6. C, D.

We found 2139 significant predictive time bins across 948 neurons that were significantly predictive if a trill call was part of a turn-taking bout in terms of median firing rate (two-sided Mann-Whitney’s U-test, p < 0.05). Only 1098 neurons were considered in these trill call analyses by the minimal session selection criteria, which was used equally in all predictive time bin analyses. We found 1545 predictive time bins across 417 neurons that were significantly predictive if a twitter call was part of a turn-taking bout (two-sided Mann-Whitney’s U-test, p < 0.05). Only 682 neurons were considered in these twitter call analyses by the same minimal session selection criteria.

*Supplementary references:* References 51 to 56 are cited only in the supplementary material.

## Supplementary Figures

**Supplementary Fig. 1.**
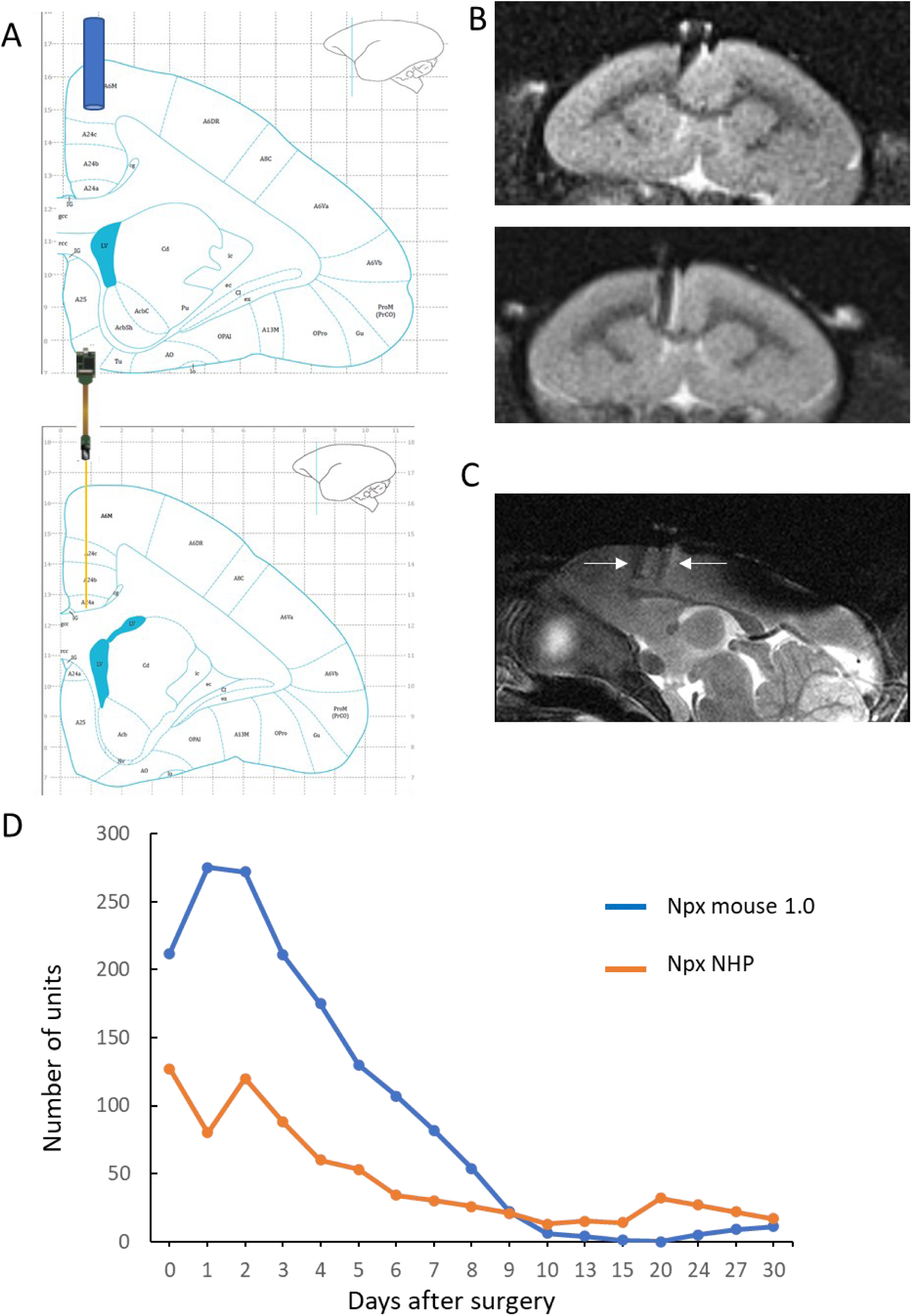
**A)** Coronal section from the Paxinos atlas showing the location of brush array (top) cannula insertion and Neuropixel (bottom) implantation site. **B)** Coronal MRI planes of monkey S before the first recording session (top) and after the last one (bottom), showing the trajectory of the 64 channels brush array through area 24. **C)** Sagittal MRI plane of monkey G showing (white arrows) the two 32 channels brush arrays implantation sites. **D)** Number of units (total = 2355) recorded per day with the 2 Neuropixel probes implanted in monkey L. Only sessions d4 to d9 were used for data analysis.

**Supplementary Fig. 2.**
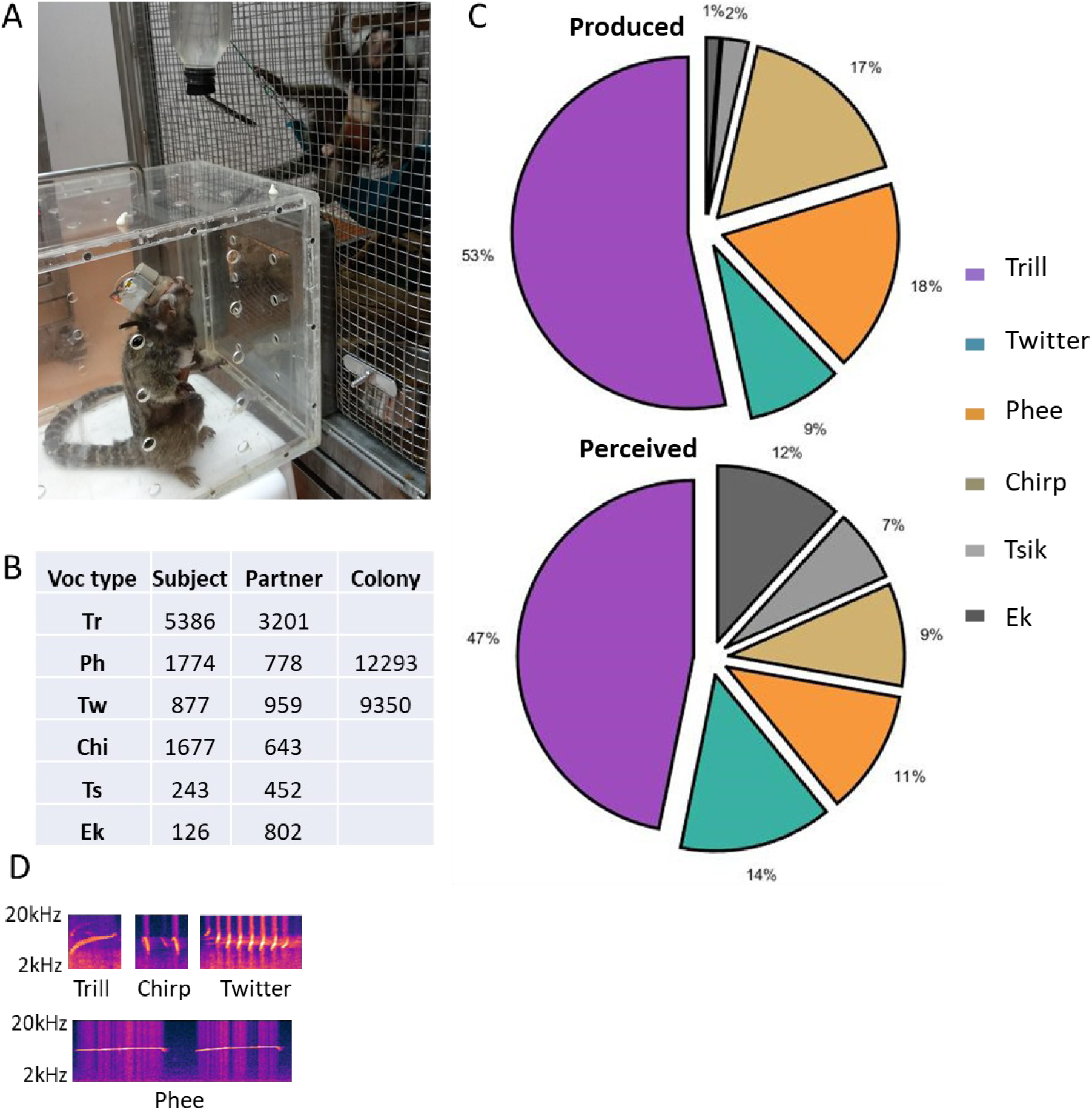
**A)** Picture taken during a recording session. **B)** Table summarizing the number of each call type per caller identity. **C)** Pie charts representing call type proportions for the subject (top) and the partner (bottom). Note that most of the perceived Tsiks and Eks came from a few specific sessions and could not be analyzed. **D)** Examples of spectrograms of recorded vocalizations.

**Supplementary Fig. 3.**
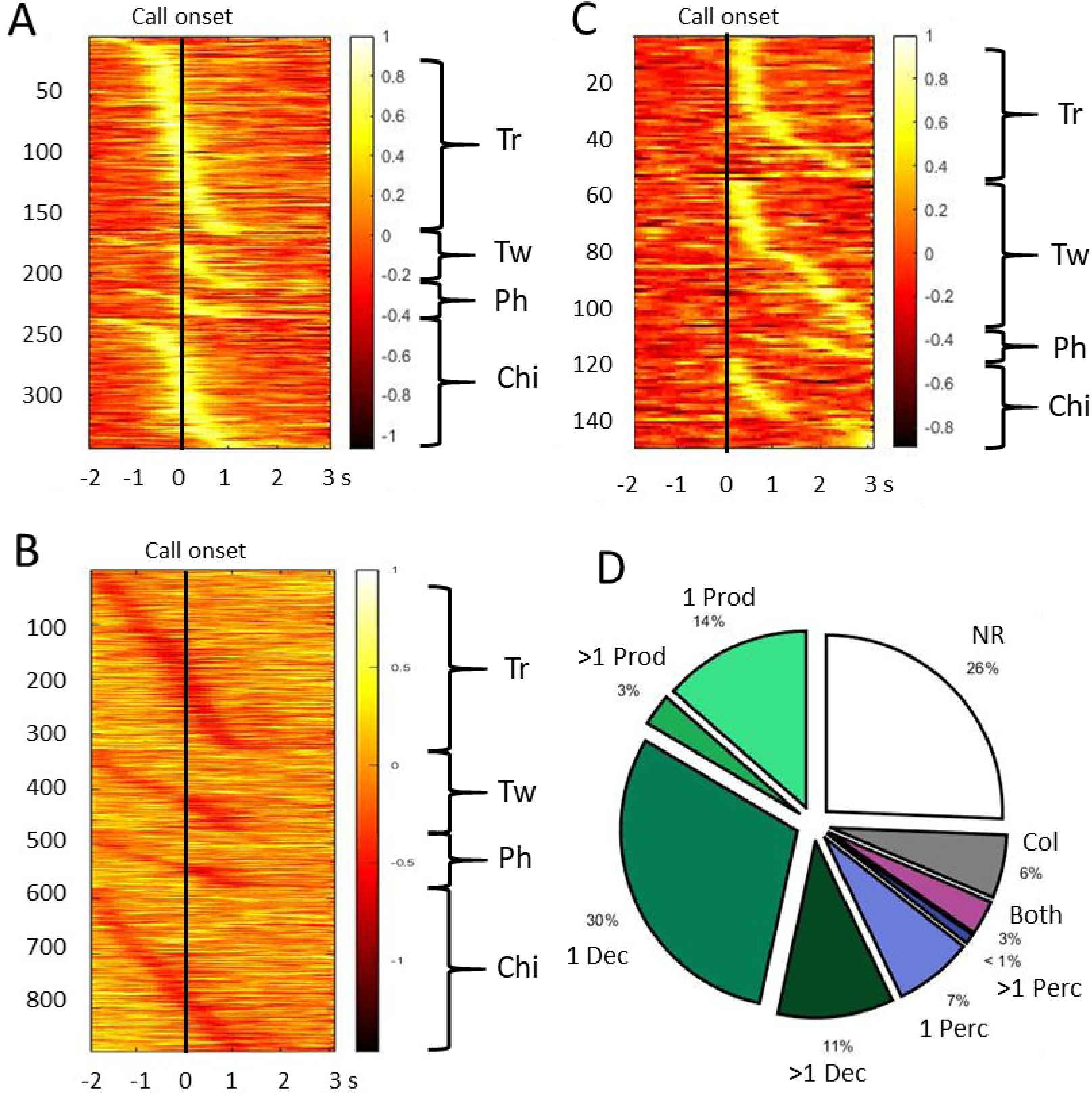
**A)** Normalized activity of neurons with significant increased response to vocalization production, per call type. **B)** Normalized activity of neurons with significant decreased response to vocalization production, per call type. **C)** Normalized activity of neurons with significant increased response to partner vocalization perception, per call type. **D)** Pie chart representing the neurons (n = 1599) responses: 217 neurons with increased activity for 1 type of vocalization production, 47 neurons with increased activity for several types of vocalization production, 482 neurons with decreased activity for 1 type of vocalization production, 170 neurons with decreased activity for several types of vocalization production, 116 neurons with increased activity for 1 type of partner vocalization perception, 15 neurons with increased activity for several types of partner vocalization perception, 93 neurons with increased activity for 1 type of colony vocalization perception, 51 neurons with increased activity for both vocalization production and partner vocalization perception, 408 neurons with no significant responses.

**Supplementary Fig. 4.**
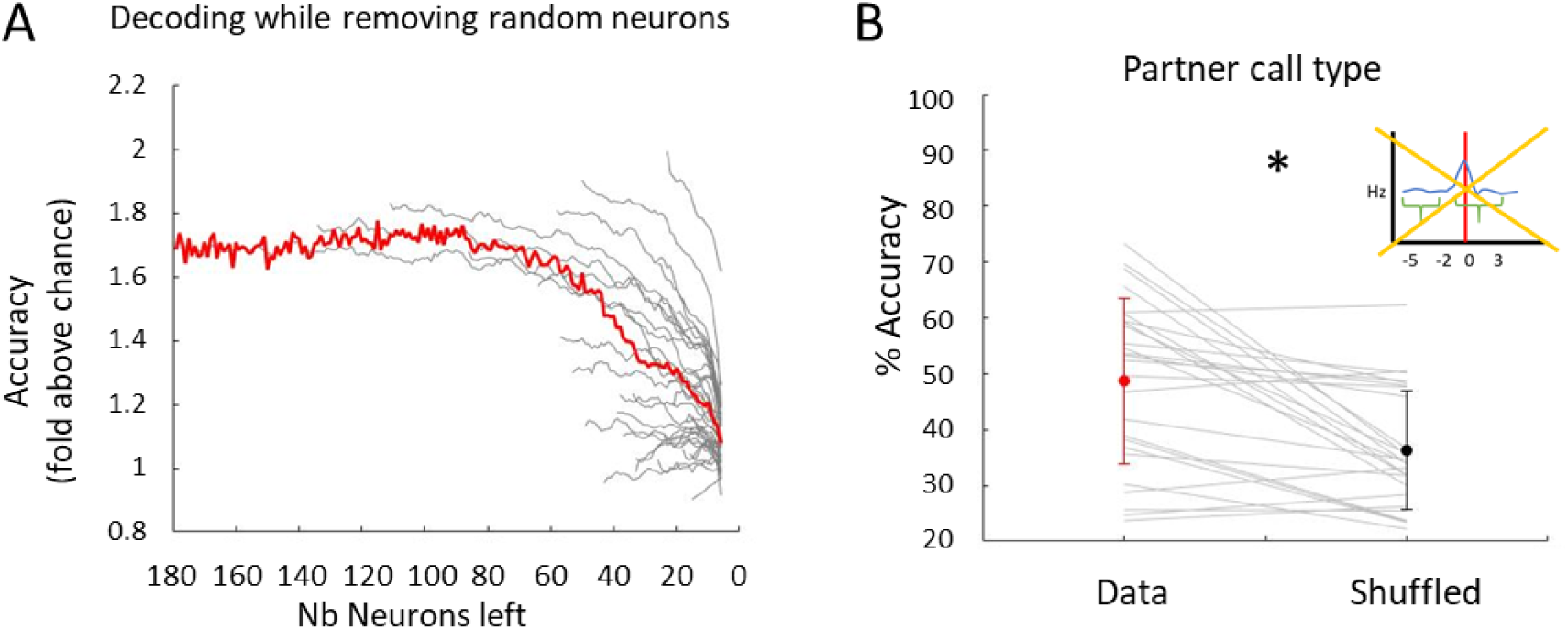
**A)** Decoding partner call type accuracy for each session (gray lines) depending on the number of neurons removed, red line is the average. **B)** Percentage accuracy of the neuronal multiclass decoder for perceived call type for real data and data with shuffled labels, after removing neurons with significant activity increase around call onset (Wilcoxon signed rank test, z = 3.65, p < 0.05). Performance was not significantly different than when significant neurons were included (Mann-Whitney U test, U = 0.47, p > 0.05). Error bars indicate s.e.m. (standard error to the mean).

**Supplementary Fig. 5.**
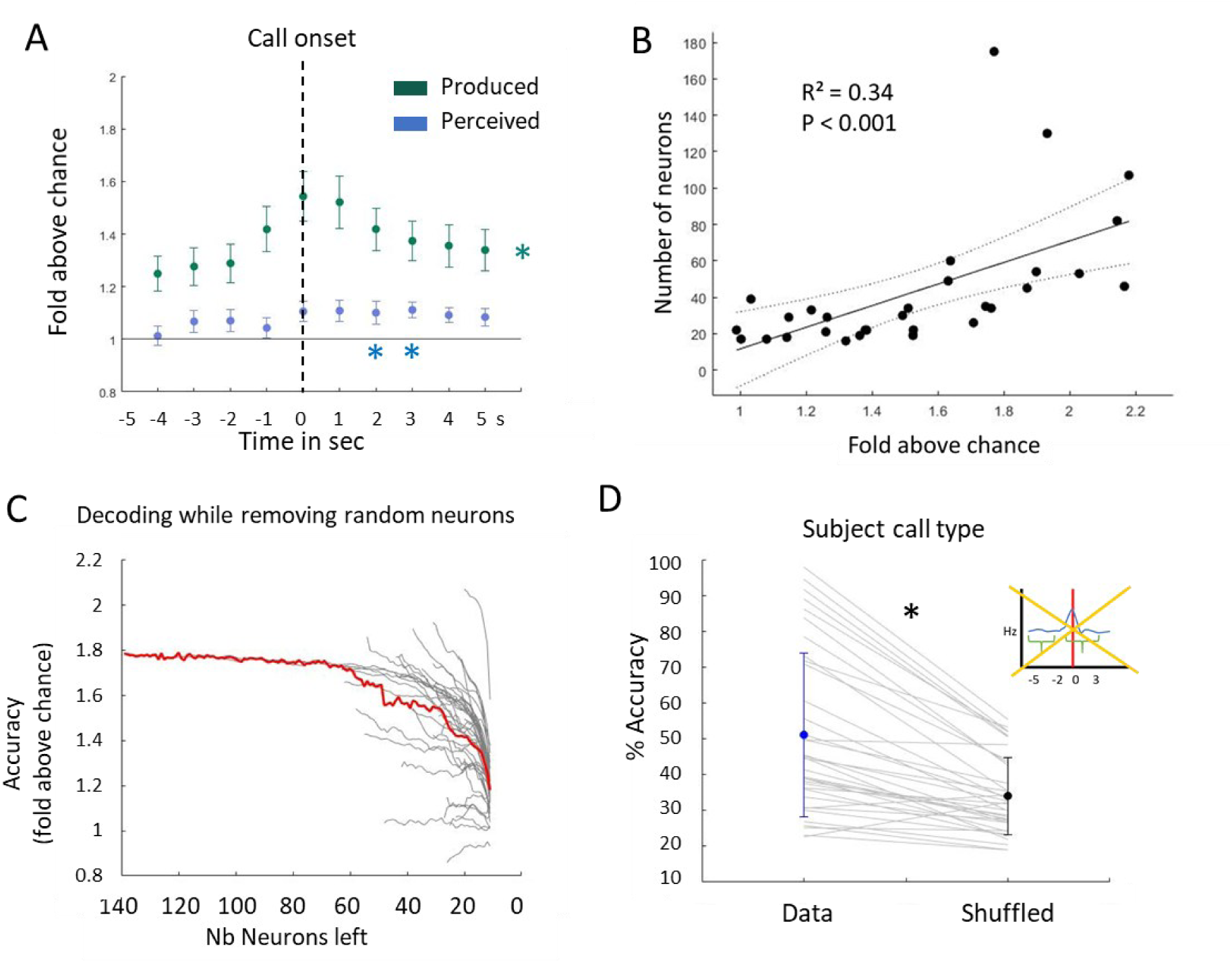
**A)** Decoding accuracy expressed in fold above chance for produced and perceived vocalizations. * indicate significant decoding compared to chance (Wilcoxon sign rank test, p < 0.05, Bonferroni corrected). **B)** Correlation between the number of neurons and the decoding quality. **C)** Decoding produced call type performance for each session (gray lines) depending on the number of neurons removed, red line is the average. **D)** Percentage accuracy of the neuronal multiclass decoder for produced call type for real data and data with shuffled labels, after removing neurons with significant activity increase around call onset (Wilcoxon signed rank test, z = 4.95, p < 0.05). Performance was not significantly different than when significant neurons were included (Mann-Whitney U test, U = 1,61, p > 0.05). Error bars indicate s.e.m. (standard error to the mean).

**Supplementary Fig. 6.**
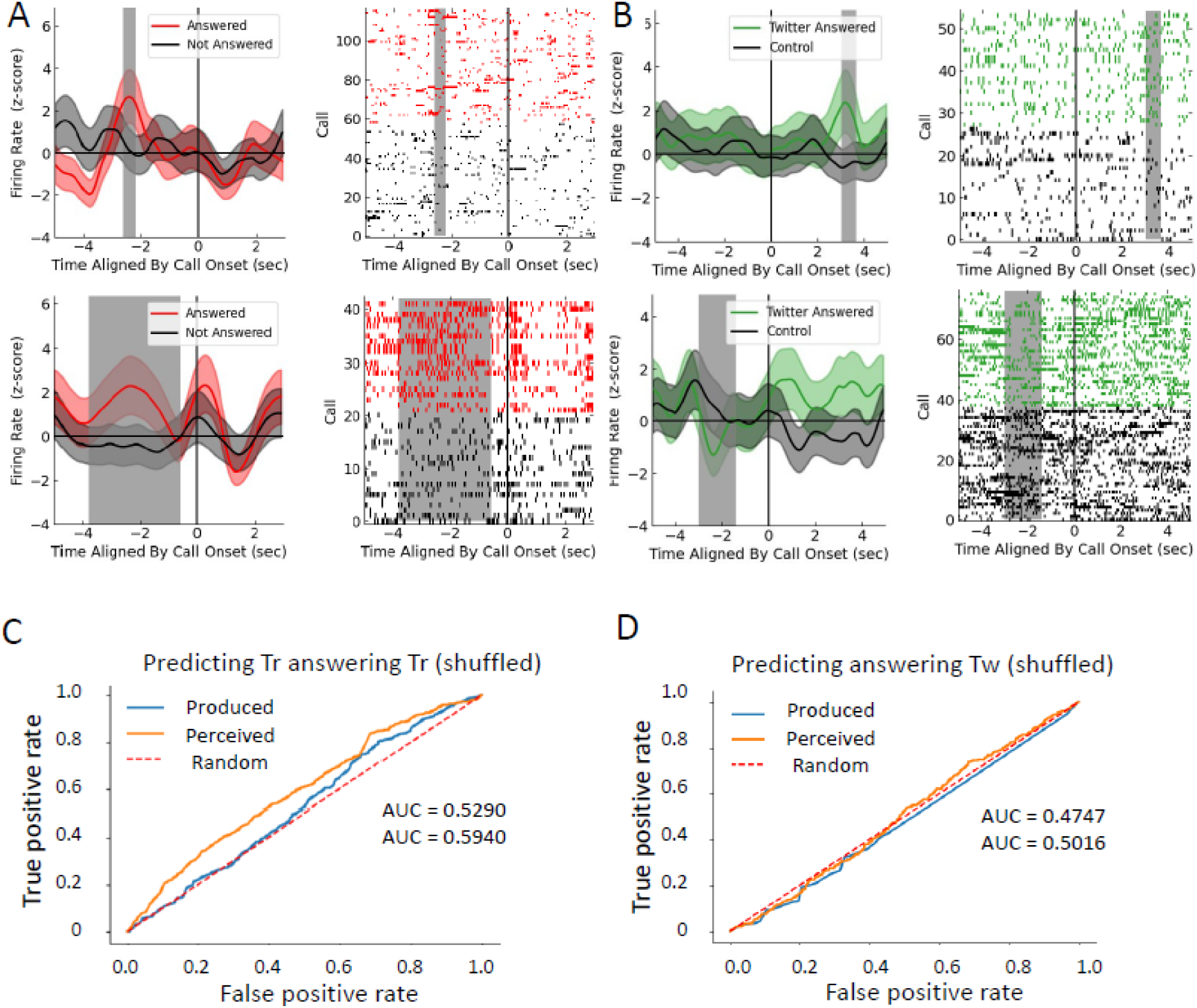
**A)** PSTH examples and spike rasters of significant time bins predicting whether a trill call was part of an exchange (within 5 sec of a trill from the other monkey). **B)** PSTH examples and spike rasters of significant time bins predicting whether a twitter call was part of an exchange (within 5 sec of a twitter from the other monkey). **C-D)** Receiver Operator Characteristic (ROC) traces for our neural decoder trained on predictive time bins and tested on data with shuffled labels representing whether trills answered trills with the subject’s partner in panel C, the answering of twitter calls with the colony in panel D. Blue solid lines represent calls initially produced by the subject while orange solid lines represent calls initially perceived by the subject. Random chance is represented by the red dashed line as a guide. The area under the curve (AUC) quantifies the success of the neural decoder as our outcome measure.

## References

1. McDermott, J. H. The cocktail party problem. Curr. Biol. CB 19, R1024–1027 (2009).

2. Mesgarani, N. & Chang, E. F. Selective cortical representation of attended speaker in multi-talker speech perception. Nature 485, 233–236 (2012).

3. Bee, M. A. & Micheyl, C. The cocktail party problem: what is it? How can it be solved? And why should animal behaviorists study it? J. Comp. Psychol. Wash. DC 1983 122, 235–251 (2008).

4. O’Sullivan, J. et al. Hierarchical Encoding of Attended Auditory Objects in Multi-talker Speech Perception. Neuron 104, 1195–1209.e3 (2019).

5. Nelken, I., Bizley, J., Shamma, S. A. & Wang, X. Auditory cortical processing in real-world listening: the auditory system going real. J. Neurosci. Off. J. Soc. Neurosci. 34, 15135–15138 (2014).

6. Joshi, N. et al. Temporal coherence shapes cortical responses to speech mixtures in a ferret cocktail party. *Commun*. Biol. 7, 1392 (2024).

7. Klein, J. T., Shepherd, S. V. & Platt, M. L. Social attention and the brain. Curr. Biol. CB 19, R958–962 (2009).

8. Jürgens, U. Neural pathways underlying vocal control. Neurosci. Biobehav. Rev. 26, 235–258 (2002).

9. Jafari, A. et al. A vocalization-processing network in marmosets. Cell Rep. 42, 112526 (2023).

10. Putnam, P. T. & Chang, S. W. C. Social processing by the primate medial frontal cortex. Int. Rev. Neurobiol. 158, 213–248 (2021).

11. Nieder, A. & Mooney, R. The neurobiology of innate, volitional and learned vocalizations in mammals and birds. Philos. Trans. R. Soc. Lond. B. Biol. Sci. 375, 20190054 (2020).

12. Burkart, J. M. et al. A convergent interaction engine: vocal communication among marmoset monkeys. Philos. Trans. R. Soc. Lond. B. Biol. Sci. 377, 20210098 (2022).

13. Miller, C. T. et al. Natural behavior is the language of the brain. Curr. Biol. CB 32, R482–R493 (2022).

14. Bezerra, B. M. & Souto, A. Structure and Usage of the Vocal Repertoire of Callithrix jacchus. Int. J. Primatol. 29, 671–701 (2008).

15. Digby, L. J. & Barreto, C. E. Social organization in a wild population of Callithrix jacchus. I. Group composition and dynamics. Folia Primatol. Int. J. Primatol. 61, 123–134 (1993).

16. Lazaro-Perea, C. Intergroup interactions in wild common marmosets, *Callithrix jacchus*: territorial defence and assessment of neighbours. Anim. Behav. 62, 11–21 (2001).

17. Miller, C. T. et al. Marmosets: A Neuroscientific Model of Human Social Behavior. Neuron 90, 219–233 (2016).

18. Burkart, J. M. & van Schaik, C. P. Marmoset prosociality is intentional. Anim. Cogn. 23, 581–594 (2020).

19. Vitale, A., Zanzoni, M., Queyras, A. & Chiarotti, F. Degree of social contact affects the emission of food calls in the common marmoset (Callithrix jacchus). Am. J. Primatol. 59, 21–28 (2003).

20. Landman, R. et al. Close-range vocal interaction in the common marmoset (Callithrix jacchus). PloS One 15, e0227392 (2020).

21. Robinson, B. W. Vocalization evoked from forebrain in Macaca mulatta. Physiol. Behav. 2, 345–354 (1967).

22. Sperli, F., Spinelli, L., Pollo, C. & Seeck, M. Contralateral smile and laughter, but no mirth, induced by electrical stimulation of the cingulate cortex. Epilepsia 47, 440–443 (2006).

23. Gavrilov, N., Hage, S. R. & Nieder, A. Functional Specialization of the Primate Frontal Lobe during Cognitive Control of Vocalizations. Cell Rep. 21, 2393–2406 (2017).

24. West, R. A. & Larson, C. R. Neurons of the anterior mesial cortex related to faciovocal activity in the awake monkey. J. Neurophysiol. 74, 1856–1869 (1995).

25. Dureux, A., Zanini, A., Trapeau, R., Belin, P. & Everling, S. Functional organization of voice patches in marmosets and cross-species comparisons with macaques and humans. Curr. Biol. CB S0960-9822(25)00874–7 (2025) doi:10.1016/j.cub.2025.07.008.

26. Bregman, A. S. Auditory scene analysis: Hearing in complex environments. (1993).

27. Petersen, S. E. & Posner, M. I. The attention system of the human brain: 20 years after. Annu. Rev. Neurosci. 35, 73–89 (2012).

28. Schneider, K. N., Sciarillo, X. A., Nudelman, J. L., Cheer, J. F. & Roesch, M. R. Anterior Cingulate Cortex Signals Attention in a Social Paradigm that Manipulates Reward and Shock. Curr. Biol. CB 30, 3724–3735.e2 (2020).

29. Benedict, R. H. B. et al. Covert auditory attention generates activation in the rostral/dorsal anterior cingulate cortex. J. Cogn. Neurosci. 14, 637–645 (2002).

30. Chen, H.-C., Kaplan, G. & Rogers, L. J. Contact calls of common marmosets (Callithrix jacchus): influence of age of caller on antiphonal calling and other vocal responses. Am. J. Primatol. 71, 165–170 (2009).

31. Li, J., Aoi, M. C. & Miller, C. T. Representing the dynamics of natural marmoset vocal behaviors in frontal cortex. Neuron 112, 3542–3550.e3 (2024).

32. Tyree, T. J., Metke, M. & Miller, C. T. Cross-modal representation of identity in the primate hippocampus. Science 382, 417–423 (2023).

33. Burgos-Robles, A., Gothard, K. M., Monfils, M. H., Morozov, A. & Vicentic, A. Conserved features of anterior cingulate networks support observational learning across species. Neurosci. Biobehav. Rev. 107, 215–228 (2019).

34. Ducret, M. et al. Medial to lateral frontal functional connectivity mapping reveals the organization of cingulate cortex. Cereb. Cortex N. Y. N 1991 34, bhae322 (2024).

35. Vogt, B. A. & Pandya, D. N. Cingulate cortex of the rhesus monkey: II. Cortical afferents. J. Comp. Neurol. 262, 271–289 (1987).

36. Bosshard, A. B. et al. Beyond bigrams: call sequencing in the common marmoset (Callithrix jacchus) vocal system. R. Soc. Open Sci. 11, 240218 (2024).

37. Oren, G. et al. Vocal labeling of others by nonhuman primates. Science 385, 996–1003 (2024).

38. Eliades, S. J. & Wang, X. Neural correlates of the lombard effect in primate auditory cortex. J. Neurosci. Off. J. Soc. Neurosci. 32, 10737–10748 (2012).

39. Tsunada, J. & Eliades, S. J. Frontal-auditory cortical interactions and sensory prediction during vocal production in marmoset monkeys. Curr. Biol. CB S0960–9822(25)00393–8 (2025) doi:10.1016/j.cub.2025.03.077.

40. Löschner, J., Pomberger, T. & Hage, S. R. Marmoset monkeys use different avoidance strategies to cope with ambient noise during vocal behavior. iScience 26, 106219 (2023).

41. Löschner, J. & Hage, S. R. Sound amongst the din: primate strategies against noise. Trends Cogn. Sci. 29, 111–113 (2025).

42. Olson, H. A., Chen, E. M., Lydic, K. O. & Saxe, R. R. Left-Hemisphere Cortical Language Regions Respond Equally to Observed Dialogue and Monologue. Neurobiol. Lang. Camb. Mass 4, 575–610 (2023).

43. Fedorenko, E., Ivanova, A. A. & Regev, T. I. The language network as a natural kind within the broader landscape of the human brain. Nat. Rev. Neurosci. 25, 289–312 (2024).

44. Ferstl, E. C. & von Cramon, D. Y. What does the frontomedian cortex contribute to language processing: coherence or theory of mind? NeuroImage 17, 1599–1612 (2002).

45. Amodio, D. M. & Frith, C. D. Meeting of minds: the medial frontal cortex and social cognition. Nat. Rev. Neurosci. 7, 268–277 (2006).

46. Wittmann, M. K., Lockwood, P. L. & Rushworth, M. F. S. Neural Mechanisms of Social Cognition in Primates. Annu. Rev. Neurosci. 41, 99–118 (2018).

47. Barris, R. W. & Schuman, H. R. [Bilateral anterior cingulate gyrus lesions; syndrome of the anterior cingulate gyri]. Neurology 3, 44–52 (1953).

48. Clairis, N. & Lopez-Persem, A. Debates on the dorsomedial prefrontal/dorsal anterior cingulate cortex: insights for future research. Brain J. Neurol. 146, 4826–4844 (2023).

49. Kolling, N., Behrens, T. E., Mars, R. B. & Rushworth, M. F. Neural Mechanisms of Foraging. Science 336, 95 (2012).

50. Abrams, D. A., Mistry, P. K., Baker, A. E., Padmanabhan, A. & Menon, V. A Neurodevelopmental Shift in Reward Circuitry from Mother’s to Nonfamilial Voices in Adolescence. J. Neurosci. Off. J. Soc. Neurosci. 42, 4164–4173 (2022).

51. Paxinos, G., Watson, C., Petrides, M., Rosa, M. & Tokuno, H. The Marmoset Brain in Stereotaxic Coordinates. (Elsevier Academic Press, 2012).

52. McMahon, D. B. T., Bondar, I. V., Afuwape, O. A. T., Ide, D. C. & Leopold, D. A. One month in the life of a neuron: longitudinal single-unit electrophysiology in the monkey visual system. J. Neurophysiol. 112, 1748–1762 (2014).

53. Oikarinen, T. et al. Deep convolutional network for animal sound classification and source attribution using dual audio recordings. J. Acoust. Soc. Am. 145, 654 (2019).

54. Pachitariu, M., Steinmetz, N., Kadir, S., Carandini, M. & D, H. K. Kilosort: realtime spike-sorting for extracellular electrophysiology with hundreds of channels. 061481 Preprint at 10.1101/061481 (2016).

55. Tyree, T. J. Applications of Mathematical Physics to Quantitative Biology. (2023).

56. Chen, T. & Guestrin, C. XGBoost: A Scalable Tree Boosting System. in 785–794 (2016). doi:10.1145/2939672.2939785.

57. Jovanovic, V., Fishbein, A. R., de la Mothe, L., Lee, K.-F. & Miller, C. T. Behavioral context affects social signal representations within single primate prefrontal cortex neurons. Neuron S0896–6273(22)00059–9 (2022) doi:10.1016/j.neuron.2022.01.020.

